# ProTargetMiner: A proteome signature library of anticancer molecules for functional discovery

**DOI:** 10.1101/421115

**Authors:** Amir Ata Saei, Alexey Chernobrovkin, Pierre Sabatier, Bo Zhang, Christian Beusch, Ülkü Güler Tokat, Massimiliano Gaetani, Ákos Végvári, Roman A. Zubarev

## Abstract

We present a publicly available, expandable proteome signature library of anticancer molecules in A549 adenocarcinoma cells. Based on 287 proteomes affected by 56 drugs, the main dataset contains 7,328 proteins and 1,307,859 refined protein-drug pairs. By employing the specificity concept in partial least square modeling, deconvolution of drug targets and mechanistic proteins is achieved for most compounds, including some kinase inhibitors. We built the first protein co-regulation database that takes into account both protein expression and degradation. A surprising number of strong anti-correlations is found, underscoring the importance of protein repression in cell regulation. Our analysis uncovered a group of proteins with extremely steady expression which are likely essential for core cellular functions. These findings bring about deeper understanding of cell mechanics. Extension of the dataset to novel compounds will facilitate drug design. The introduced specificity concept and modeling scheme are beneficial in other analysis types as well.

**Statement of Significance:** ProTargetMiner is the first of its kind library of proteome responses of human cancer cells to anticancer molecules. This expandable resource facilitates the deconvolution of drug targets, action mechanisms, and cellular effects. It reveals death modalities, uncovers protein co-regulation and anti-correlation networks and defines the “untouchable” proteome essential for core cellular functionalities.

## Introduction

Deciphering the targets, action mechanisms, resistance factors, and the modalities of cancer cell death for novel compounds, especially for those derived from phenotypic screenings, are all challenging tasks in drug discovery. These tasks can be addressed by connecting the affected cellular phenotypes to small molecules by so-called connectivity maps (1-3). Such an approach explores the similarities of the cell response signatures produced by the compound of interest to other responses in the database. However, majority of studies published so far are based on gene expression (mRNA) profiles. Since proteins are the targets of most drugs, proteome responses can be more specific to drug action. One recent effort has focused on building a connectivity map based on phosphoproteomic and chromatin signatures, measuring the abundances of 100 phosphopeptides and 59 histone modifications for treatments with 90 drugs (4). There is no reason why protein abundances cannot serve a basis for connectivity maps; after all, biological systems are defined by their proteome status. Also, protein abundances are in general as much determined by expression as by degradation (5), both reflected uniquely in proteomics data. In several studies, no strong correlation between mRNA levels and protein concentrations has been found even at the steady state (6). In dynamic situations where degradation processes play an important role, such as programmed cell death (7), the relationship between the transcriptome and proteome should become even less direct.

Here we use chemical proteomics to study the relationship between the anticancer drug molecules and the dying cell phenotype induced by these molecules. Chemical proteomics has traditionally been defined as the use of small molecules (which are considered known entities) in studying the unknown functions of proteins (8). Recently, chemical proteomics began to designate also the opposite approach, in which proteome analysis is applied to studying functions of small molecules (9).

We have previously shown that, when sensitive cell lines are treated with the compound of interest, drug targets are consistently found among the most regulated proteins and mapping these proteins on protein networks can reveal the drug action mechanism. This observation served as a basis for a new chemical proteomics method called Functional Identification of Target by Expression Proteomics (FITExP) (10). In many cases, the affected target and other mechanism-related proteins are found upregulated. This can be explained by a feedback effect when inhibition of certain proteins activates their (over)production. The alternative effect, involving target protein down-regulation, can be caused by protein translocation followed by proteolysis (11).

To increase the specificity of analysis for a given compound, a panel of known drugs is added to the experiment. The *specificity* parameter reflects the protein regulation in the treatment by a given compound versus all other compounds. FITExP could successfully identify the targets of several chemotherapeutics (10), and probe the targets and mechanisms of metallodrugs (12) and even toxic nanoparticles (13). We have also shown that combining the proteomic data from treated matrix-attached cells with matrix-detached cells can improve the deconvolution of drug targets and action mechanisms, and identify proteins involved in cell life and death decisions (14). Achieving a high level of specificity in analysis usually requires the use of several compounds and cell lines. We hypothesized that the equivalent increase in specificity can be obtained with a single cell line when a multitude of “contrasting” drugs is used.

While specifically regulated proteins help to identify the drug targets and the action mechanisms, mapping the whole proteome, and even the set of ≥1000 most abundant proteins, may help to determine the cell death modality (15). Assuming that the hypothesis on the molecule-specific nature of the obtained dataset holds, interrogating a cell line with an extensive (>50 molecules) drug panel was expected to probe most cell death pathways. Each pathway can be represented by a cell death trajectory in a space encompassing all possible cell states. Therefore, we could hope to determine the number of orthogonal death modalities – the subject that has generated a significant debate and even controversy in the cell death community (16).

We define a death trajectory caused by a toxic agent, to be a track in the proteome space, passing from a normal living state that the untreated cells occupy and explore during cell division, circadian and metabolic cycles, to a particular death state. The proteome space and death trajectories are schematically shown in Fig. 1. The first crossing beyond the normal state defines the molecular action mechanism of the toxic agent.

**Figure 1.**
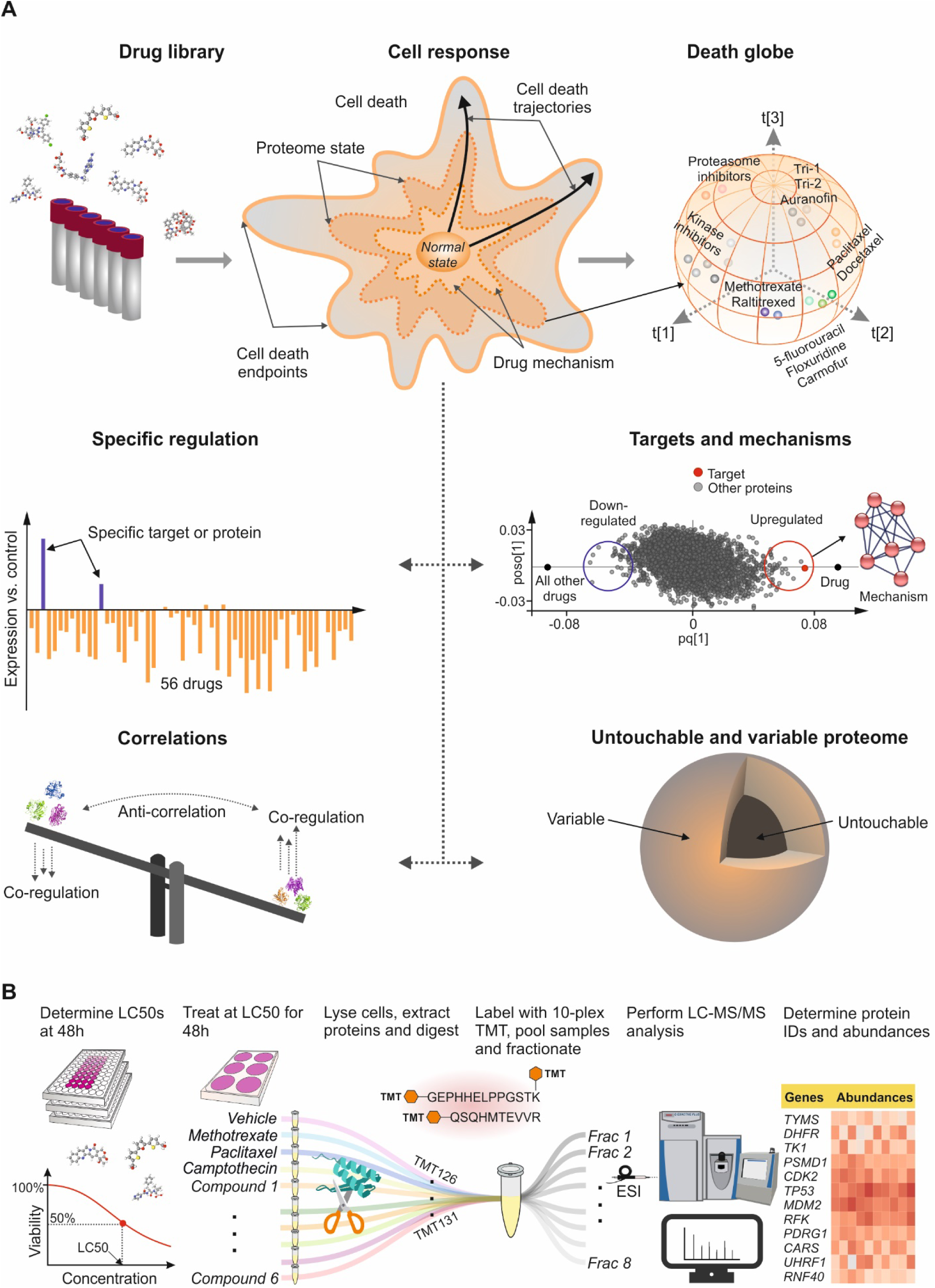
Objectives and workflow of ProTargetMiner. **A,** Interrogating a cell line with an extensive drug panel will probe most cell death pathways, each represented by a different cell death trajectory, which will allow us to build a global cell death map (multidimensional “Death globe”), for clustering of compounds. The extensive proteome dataset will also provide specific information on drug targets, mechanistic proteins, and other specifically regulated proteins. The proteome perturbation data will provide novel protein co-regulation and anti-correlation links, as well as define untouchable and variable proteomes. **B,** Workflow: determination of LC50 values for the compound library; cell treatment with 56 compounds and sample preparation for shotgun proteomics; sample multiplexing including control and standard treatments (methotrexate, paclitaxel and camptothecin); lysis, digestion and labeling with TMT-10plex; combining the 10-plexed samples and fractionation of the pooled sample to increase the proteome coverage; analysis of individual fractions by LC-MS/MS; protein identification and quantification; data post-processing.

We selected A549 human adenocarcinoma cells originating from lung cancer as a model system, because it is well covered in the literature, and it showed the highest sensitivity to the compounds among other tested cell lines (MCF-7 and RKO) in a pilot experiment. Viability measurements were performed for 118 clinical molecules cherry-picked for cancer from Selleckchem FDA-approved drug library and several experimental compounds with unknown targets. A collection of 56 compounds were chosen to treat the cells at LC50 concentrations for 48 h (selection criterion was the induction of 50% cytotoxicity in 48h at concentrations below 50 *µ*M). With the biological effect (cell death) being of the same magnitude, the differences in the proteome states could be attributable to the differences in targets, action mechanisms as well as modalities of cell death and survival. The selected compounds belong, according to available information, to 18 different classes with versatile targets and mechanisms, spanning 112 known drug targets curated from DrugBank (https://www.drugbank.ca/) in September 2018. These drugs and their known targets are listed in Supplementary Table S1. As standard drugs used for quality control, methotrexate, paclitaxel, and camptothecin were chosen and included in each TMT10 multiplexed proteomics experiment (labeling information for each experiment is given in Supplementary Table S2).

The overview of the project’s objectives and the workflow is given in Fig. 1. Besides mapping the orthogonal death pathways, the obtained dataset allowed us to deconvolve compound targets and action mechanisms. We could also probe protein co-regulation and anti-correlation networks, the untouchable proteome that is (almost) unchanged in all perturbations, as well as the variable proteome that exhibits large drug-to-drug variations. Here we present this dataset, called ProTargetMiner, as well as our most significant findings so far from its interrogation.

## Results

### Overview of the Proteomics data

Overall, for the original dataset, 287 cell lysates were prepared and 229 LC-MS/MS analyses were performed. In total, 144,075 peptides attributed to 7,328 proteins were quantified in all experiments, with at least 2 unique peptides per protein. After selecting only proteins quantified in at least one replicate in each treatment, the list was reduced to 4,557 proteins (Supplementary Table S3) that were used in all subsequent analyses. The final resource therefore, contains 1,307,859 (287×4557) filtered high quality drug-protein data points.

### Number of independent dimensions

Using factor analysis of the whole dataset, we identified at least 11 independent dimensions. Supplementary Table S4 lists the contribution of all proteins to each dimension. The most contributing proteins to the first dimension were cyclin-dependent kinase inhibitor 1 (*CDKN1A*) and *PCNA*-associated factor (*PCLAF*), with opposite signs. While *CDKN1A* is involved in *p53* mediated inhibition of cellular proliferation by blocking cell cycle progression, *PCLAF* is a cell cycle-regulated substrate that acts as a regulator of DNA repair during replication. The top 30 proteins of the first three dimensions map best to “p53 signaling pathway and cell cycle”, “focal adhesion and angiogenesis” and “chromatin assembly and fatty acid metabolism”, respectively. Some of the dimensions clearly correspond to classical cell death modalities, such as p53-dependent apoptosis (1^st^ dimension), autophagy (4^th^ dimension), and macromitophagy (6^th^ dimension), while other dimensions were harder to ascribe to known modalities. Associating top proteins defining these dimensions with cell death modalities is a subject of future research. Supplementary Table S5 lists the top two KEGG and biological pathways for 30 most-contributing proteins for all 11 dimensions.

### Drugs with similar mechanisms induce similar proteome changes

We employed a nonlinear dimension reduction method t-SNE that is widely used for projection of molecular signatures in transcriptomics, to reduce the proteomic space to three dimensions (17). As a result, we obtained the “Death globe”, on which all drug-induced proteome signatures are mapped as points. We used the proximity of these points in the 3D t-SNE projection to evaluate the similarity of the drug-induced signatures. As expected, drugs with similar mechanisms (*e.g.*, tubulin depolymerization inhibitors paclitaxel and docetaxel, pyrimidine analogues 5-fluorouracil, floxuridine and carmofur, thioredoxin reductase inhibitors auranofin, TRi-1 and TRi-2 (18) and topoisomerase inhibitors camptothecin, topotecan and irinotecan) were proximate on the t-SNE plot, confirming that the Death globe approach can be used for evaluating the similarities between drug action mechanisms. Similar results were obtained with a 2D t-SNE plot (“Death map”, Fig. 2A). We also employed a conventional correlation-based hierarchical clustering analysis to confirm the t-SNE findings (Fig. 2B).

**Figure 2.**
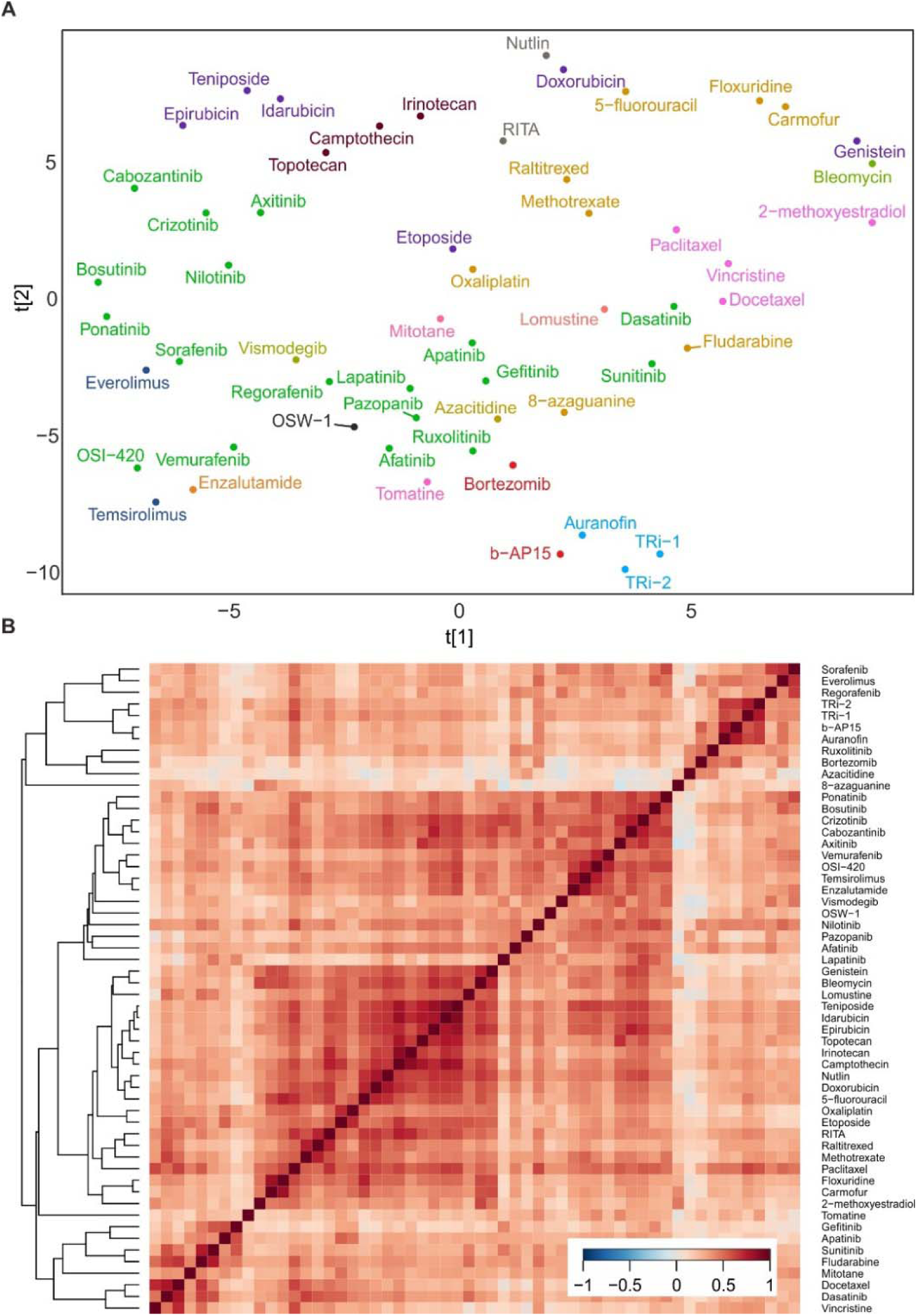
Drugs with similar mechanisms induce similar proteome changes. **A,** The compounds with similar mechanisms (with the same colors) were found to be proximate on the 2D t-SNE plot (“Death map”). **B,** Correlation based heatmap with the hierarchical clustering analysis was used to support the findings from the t-SNE.

We found tomatine to be a gross outlier in the t-SNE and PCA plots and thus excluded it from the subsequent analyses. Tomatine is likely to act via proteasome inhibition (19), along with unspecific membrane damage (20). These mechanisms may explain the extraordinary changes induced by tomatine in the cell proteome.

### ProTargetMiner enables functional discovery

Protein regulation is usually defined as a ratio of the protein abundances in the cells incubated with a drug and vehicle control. However, regulation is not a parameter specific enough in drug target deconvolution because of the proteins involved in generic cell responses to toxic agents (*e.g.*, death or survival pathways). Instead, we introduced “specific regulation”, which was defined as the ratio of the regulation to a particular drug to the median regulation in all other drugs of the panel (10). Here, we also employ a sophisticated supervised classification method called orthogonal partial least square discriminant analysis (OPLS-DA), for the discovery of specifically regulated proteins (21). With this approach, different models can be easily built, contrasting a given compound, or a group of compounds, against either all other molecules or a selected subset. In these models, the proteins specifically up- or down-regulated in response to a given treatment are found on the opposite sides of the loading plot, with each protein represented by a dot (Fig. 3A). The position on the y-axis reflects the strength of the orthogonal components, and best target candidates are thus located near the x-axis extremities.

**Figure 3.**
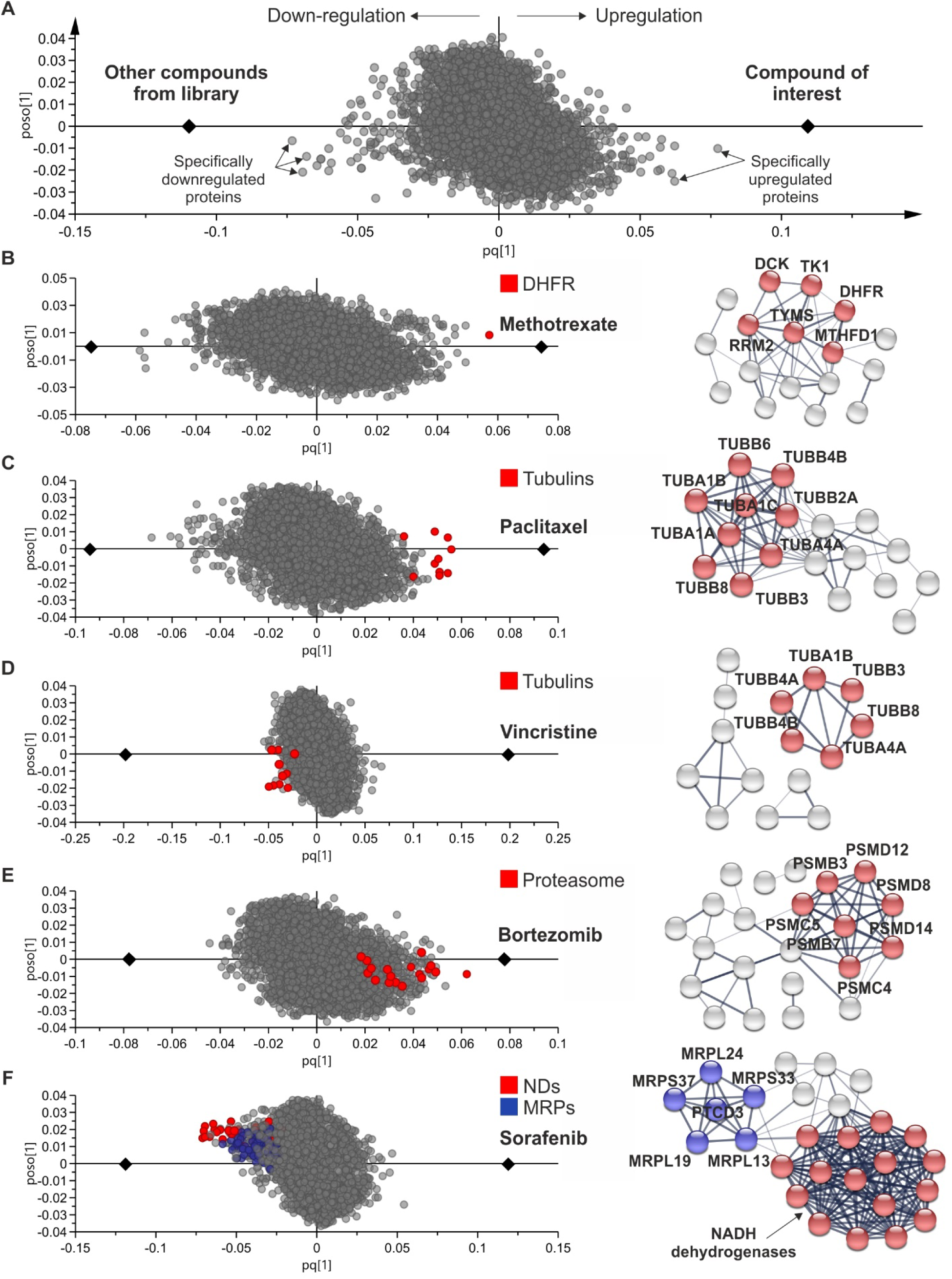
ProTargetMiner highlights the targets, action mechanisms as well as affected cellular complexes. **A,** A generalized OPLS-DA model contrasting a given compound against all others. **B-F,** Representative OPLS-DA models for five compounds. The right panel shows the mechanistically relevant pathway enrichment for the 30 most specifically up- or down-regulated proteins: KEGG pathways for methotrexate - “one carbon pool by folate” (p<0.001) and “pyrimidine metabolism” (p<0.002); paclitaxel - “tubulin” (p<2E-16); vincristine - “tubulin” (p<4E-08); bortezomib - “proteasome” (p<7E-11). GO terms for sorafenib - “NADH dehydrogenase activity” (in red, p<4E-33) and “mitochondrial translation” (in blue, p<5E-07). MRP = mitochondrial ribosomal proteins, ND = NADH dehydrogenase.

As representative examples of drug target deconvolution, OPLS models for methotrexate, paclitaxel, and vincristine are shown in Fig. 3B-D, respectively. The methotrexate target *DHFR* is convincingly identified. Tubulins are found to be most specifically up-regulated proteins for paclitaxel and down-regulated for vincristine. These two drugs affect tubulin depolymerization and polymerization, respectively. Network analysis of the specifically regulated proteins on either side of the model highlights the compounds’ mode of action (right panel in Fig. 3).

ProTargetMiner can also reveal a compound effect on protein complexes and organelles. The proteasome inhibitor Bortezomib demonstrates enrichment of the proteasome subunits among specifically up-regulated proteins (Fig. 3E). The sorafenib model shows the specific down-regulation of NADH dehydrogenases and mitochondrial ribosomal proteins (Fig. 3F). This latter finding is in line with the earlier report for human neuroblastoma cells (22); thus the ProTargetMiner results can be cautiously generalized to other cells.

As another example, pyrimidine analogues floxuridine and carmofur down-regulated ribosomal proteins similar to 5-fluorouracil (23) (Supplementary Fig. S1A). A similar effect on ribosome was also observed for the alkylating agent oxaliplatin (Supplementary Fig. S1A). A very recent finding shows that oxaliplatin kills cells by inducing ribosome biogenesis stress, unlike its platinum analogues (24).

In Supplementary Table S6, the specificity (extracted from OPLS models) of each protein in response to each compound (against all other compounds) is provided.

### ProTargetMiner analysis of kinase inhibitors

One could suspect that ProTargetMiner would be less sensitive to kinase inhibitors because this analysis doesn’t include assessment of the phosphorylation sites, while there are fewer reasons to expect for the kinase abundance to be affected by its inhibition than for metabolite-processing proteins or generally other targets. To investigate the efficiency of ProTargetMiner in the analysis of kinase inhibitors, we built a PCA plot solely based on the expression of 68 kinases with no missing values in the ProTargetMiner dataset. Both principal components 1 (18.8%) and 2 (10.5%) could crudely separate kinase inhibitors from other compounds, and especially topoisomerase and proteasome inhibitors (Supplementary Fig. S1B). Thus, with certain limitations, ProTargetMiner is also applicable to kinase inhibitors.

ProTargetMiner yields unexpected findings for kinase inhibitors that could be mechanistically relevant. As an example, gefitinib and lapatinib up-regulated proteins involved in lipid synthesis and cholesterol metabolism, indicating potential inhibition of these pathways (Supplementary Fig. S2A). This finding may explain the reduction of serum cholesterol level in patients treated with gefitinib (25). Other kinase inhibitors in our library which affected proteins involved in lipid synthesis and cholesterol metabolism were bosutinib, crizotinib, sunitinib, and cabozantinib. The specific regulation of the three up-regulated proteins involved in cholesterol metabolism in response to the mentioned compounds is shown in Supplementary Fig. S2B.

Biochemical pathways affected by compounds are related not only to death pathways but also to cell survival (14). Therefore, the specifically regulated proteins could be potentially linked to drug resistance. For example, *EGFR* was specifically upregulated in the sorafenib model (Supplementary Fig. S2C) and is known to be involved in resistance to this drug (26). Another kinase upregulated in response to sorafenib (and regorafenib) was *AXL* (Supplementary Fig. S2C-D). *AXL* is a receptor tyrosine kinase regulating many aspects of cell proliferation and survival, and its overexpression induces resistance to *EGFR* targeted therapies (27). To investigate if *AXL* inhibition can rescue A549 cells from sorafenib and regorafenib, we combined these molecules with TP0903, a specific *AXL* inhibitor in non-cytotoxic concentrations (<100nM). The combination treatment significantly sensitized the cells to sorafenib and regorafenib in 24 h and 48 h (Supplementary Fig. S2D).

### The degree of drug-induced proteome changes

The degree of proteome perturbation can vary likely due to a difference in the number of affected targets and pathways. To test this hypothesis, the drugs were ranked by the overall deviation of their molecular signatures compared to the untreated state (Fig. 4). Bortezomib induced the highest proteome variation, while tubulin inhibitors gave the least proteome perturbation potentially indicating the lack of off-target effects. As an alternative deviation assessment, the number of up- or down-regulated proteins was calculated for each compound at a 1.5 fold cutoff (Supplementary Fig. S3A). For proteasome inhibitors b-AP15 and especially for bortezomib, the number of significantly up-regulated proteins was much higher than down-regulated proteins (up/down ratio of 17.8 for bortezomib compared to the average of 2.9 for all other drugs). The variation induced by bortezomib was much larger than by b-AP15, likely indicating that bortezomib is affecting more targets/pathways. As another example, the proteome variation was larger for auranofin compared to other thioredoxin reductase inhibitors TRi-1 and TRi-2, confirming the recent finding that auranofin has off-targets (18).

**Figure 4.**
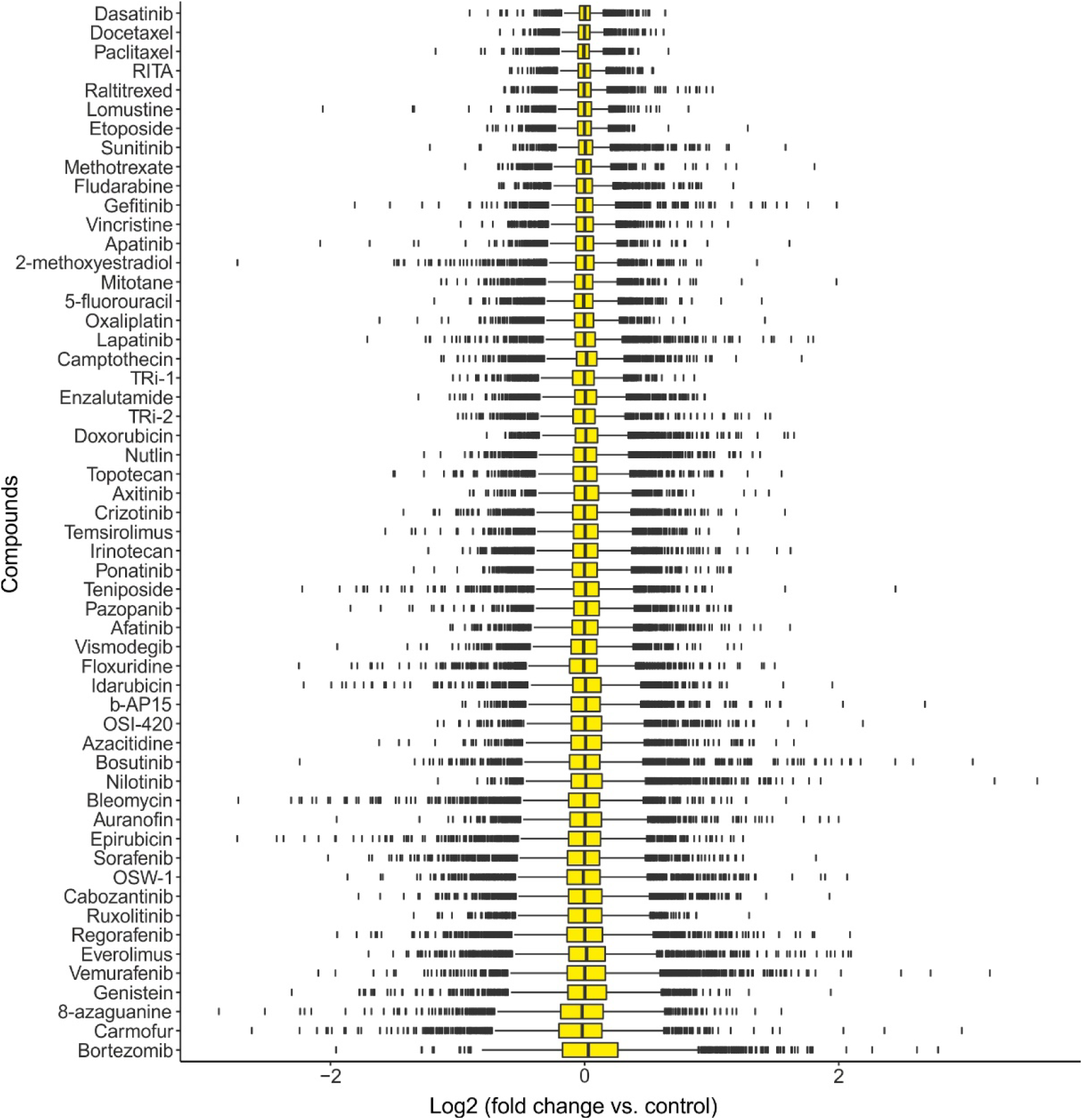
The extent of variation at the proteome level induced by each compound. Total variation of proteome changes might be indicative of compound specificity. See also Supplementary Fig. S3.

### Complex-specific effects of compounds

To investigate the drug effects on selected protein complexes, we plotted the mean specific regulations for 74 ribosomal proteins, 66 mitochondrial ribosomal proteins, and 40 proteasome subunits (Supplementary Fig. S3B-E). 5-fluorouracil, floxuridine, carmofur, and oxaliplatin showed outstanding down-regulatory effects on the ribosome (Supplementary Fig. S3B). Mitochondrial ribosome down-regulation was a specific feature for sorafenib, regorafenib, everolimus, mitotane and b-AP15 (Supplementary Supplementary Fig. S3C), most of which have documented effect on mitochondrial activity (22,28). The largest effects on proteasome were caused by bortezomib and b-AP15 (Supplementary Fig. S3D). The mean proteasome regulation was only 1.15 fold (Supplementary Fig. S3E), confirming the high specificity of the OPLS-enabled analysis and precision of measurements.

In PCA plots built using proteins with no missing values (74 ribosomal, 57 mitochondrial ribosomal and 38 proteasomal proteins), the molecules that affect the respective complexes were well separated from other compounds (Supplementary Fig. S3F-H, respectively). Interestingly, auranofin was proximal to proteasome inhibitors (Supplementary Fig. S3H), in line with a recent work implicating auranofin in inhibition of proteasomal deubiquitinase (29).

### A protein correlation database

As protein synthesis is a resource demanding cell function, protein expression is spatiotemporally controlled. Coordinated expression seems optimal for a set of genes involved in the same protein complex or biological pathway. Multiple studies have shown that co-expression can indicate functional relationships. Therefore, co-expression has been extensively exploited for deduction of gene function (1) through association analysis (30). However, co-regulation databases are mainly available based on transcripts expression (30,31), while proteomics can capture co-regulation more accurately (32). Yet to the best of our knowledge, no large-scale proteome co-regulation database has been made public so far.

ProTargetMiner provides an opportunity to build such a database of protein co-regulation. We analyzed the pairwise correlations of proteins in a 4212 x 4212 matrix (Fig. 5A). At least 11 clusters can be discerned (Fig. 5A; the constituent proteins and enriched pathways are presented in Supplementary Table S7). For example, cluster 8 contains 129 proteins, of which 72 proteins are ribosomal and 10 are involved in ribosome biogenesis.

**Figure 5.**
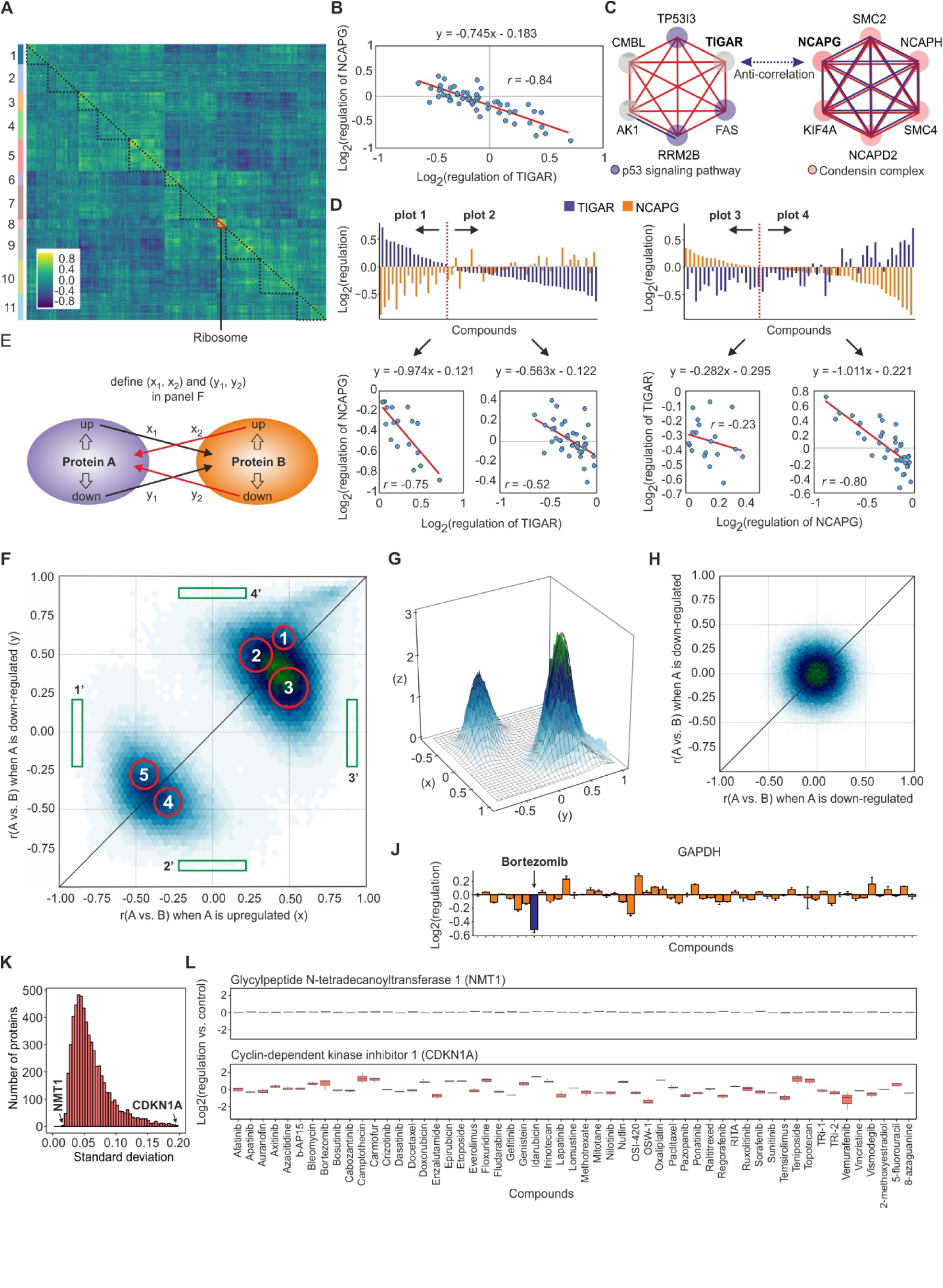
Analyzing protein pairwise correlations. **A,** The overall correlation matrix of 4,212 protein across all treatments. Vertical axis: break-down to 11 clusters. Note that the abundance of blue color (anti-correlations) is comparable with that of yellow color (positive correlations). **B,** The anti-correlation of *TIGAR* with *NCAPG*. **C,** The proteins co-regulating with *TIGAR* anti-correlate with *NCAPG* co-regulating proteins. The blue color reflects information present in StringDB and the red color reflects newly established links. **D,** The anti-correlation of up or down-regulated *TIGAR* vs. *NCAPG* (plots 1 and 2) and up-or down-regulated *NCAPG* vs. *TIGAR* (plots 3 and 4). **E,** Any two proteins give two pairs of correlations considering separately their up and down-regulation states. **F,** Map of pairwise correlations between proteins A and B, separately for up- and down-regulations of protein A. The red circles reflect the positions and volumes of 5 fitted 2D Gaussians. The green boxes encompass protein pairs that strongly correlate or anti-correlate only when protein A is up- or down-regulated. **G,** 2D Gaussian fitting is best with 5 components. **H,** Similar analysis as in F using scrambled protein abundances gives a single central spot. **J,** The expression of *GAPDH* across the treatments. **K,** Distribution of standard deviations of 6,032 protein abundances across the treatments. **L,** Representative examples of proteins showing lowest or highest expression variability across the treatments. Data are represented as mean±SD.

At FDR<0.001, a high-confidence set of 103,928 positively and 51,137 negatively correlating protein pairs were found (Supplementary Table S8), representing approximately 1% of the total of 17,740,944 pairs. The less frequency of negative compared to positive correlations is in line with a previous study (33).

Out of the 10 strongest correlating pairs (*r*>0.98), most belong to dense regions of protein interaction networks, *e.g.*, protein complexes, such as MCM, condensin, ribosome, chaperonin-containing T-complex and mitochondrial respiratory chain complex, or to the same protein superfamily, such as tubulins. Supplementary Fig. S4A maps the top 2500 co-regulated pairs, mostly originating from the same complexes or pathways, showing functionally coherent groups of genes. The 128 proteins (3% of all proteins) exhibiting no strong correlation with any other protein (Supplementary Table S8) mapped to “RNA binding” (24 proteins, p<0.017) and “catalytic activity” (52 proteins, p<0.017).

We also calculated the number of total co-regulating (and anti-correlating) partners for each protein, to reveal hub proteins (Supplementary Fig. S4B-C). Mitochondrial trifunctional enzyme subunit beta (*HADHB*) and alpha (*HADHA*) with 276 and 271 co-regulation partners were on top.

For network visualization and pathway analysis, we mapped the proteins to StringDB entities and generated a set of external payload data, which can be uploaded to the StringDB or its plugin in Cytoscape using personal configurations (nodes as well as positive and negative edges are available in Supplementary Table S9). This way, known interactions and ProTargetMiner co-regulations can be visualized on the same plot.

The anti-correlation of protein abundances is often ignored, even though negative correlations are less likely than positive correlations to arise from technically induced artifacts (33), and can reveal opposing biological processes. For example, anti-correlations can reflect active regulatory transcriptional repression, activation or even canceling of such events (34). Therefore we investigated this aspect in more detail.

Since *TP53*-inducible glycolysis and apoptosis regulator (*TIGAR*) was among the top anti-correlating proteins with many anti-correlating protein pairs, we chose to compare its expression pattern with its most anti-correlating pair *NCAPG* (*r*=−0.84) (Fig. 5B). *TIGAR* and *NCAPG*, each one together with 5 most correlating proteins, were projected to protein networks (Fig. 5C). The *TIGAR* group mapped to p53 signaling pathway (3 proteins, p<0.002), while the *NCAPG* group mapped to condensin complex (5 proteins, p< 2.6E-13) and cell division (6 proteins, p<2.98E-07). This finding is in line with p53 mediated inhibition of entry into mitosis when DNA synthesis is blocked, and with *TIGAR*’s role in p53-mediated protection from the accumulation of genomic damage (35). *CMBL* strongly co-regulated with *TIGAR* (*r*=0.89), but was not part of the p53 signaling pathway in String. However, a microarray screen found *CMBL* to be a p53-inducible protein (36), confirming the predictive power of protein co-regulations.

Analyzing in detail the *TIGAR* vs *NCAPG* dependence (Fig. 5D), we noticed that while the overall correlation coefficient was significant (r=−0.84), the slope −0.75 was far from −1, indicating an underlying complexity. A closer inspection revealed that, when *TIGAR* had an above-average regulation, its anti-correlation with *NCAPG* (*r*=−0.75, plot 1 on Fig. 5D) was stronger than when it was relatively down-regulated (*r*=−0.52, plot 2). Also, with TIGAR up-regulated, its slope with *NCAPG* was close to −1 (−0.97), as in perfect association. However, when *TIGAR* was down-regulated, the slope declined as well (−0.56). In contrast, when *NCAPG* was up-regulated, its anti-correlation with *TIGAR* was weak (r=−0.23, plot 3), but when *NCAPG* was down-regulated, it anti-correlated with *TIGAR* very strongly (*r*=−0.80, slope −1.01, plot 4). These findings showed that the anti-correlation can be bimodal, which led us to consider two pairs of correlation coefficients for every protein pair (Fig. 5E), with correlations calculated separately for above and below the median abundance of protein A. When these correlations were heat-mapped on a 2D plot with a cutoff of |*r*|>0.54 (Fig. 5F), five dense areas unexpectedly emerged (circles 1-5 in Fig. 5G) (the correlation data can be found in Supplementary Table S10). Fitting symmetric 2D Gaussian distributions confirmed the presence of 5 entities. The positions of these Gaussians coincide with the centers of red circles in Fig. 5G, and their volumes are reflected in the areas of the circles.

At first glance, the spots 2-4 look too symmetric in respect to the main diagonal, which raises a suspicion of an artifact. Indeed, if the spot produced by a single group of proteins had an elliptic rather than circular symmetry, then two symmetric Gaussians would fit to it instead of one. However, scrambled protein abundance data gave a single central spot with a circular rather than elliptic symmetry (Fig. 5H). Moreover, the spot 1 doesn’t have a symmetric counterpart. Therefore, entities 2-4 appear to be real.

Entity 1 (“highly co-regulated”) composes 12% of the total number of (anti-) correlating protein pairs, encompassing pairs with a high co-regulation (average r=0.52). The top 300 protein pairs (*r*≥0.90) mapped to ribosome, MCM and condensin complexes, chaperonin-containing T-complex, NADH dehydrogenases, pyruvate dehydrogenase, tubulin superfamily as well as some integrins and spectrins. This component is centered above the diagonal, meaning that co-regulation is higher when the proteins are down-regulated than when they are upregulated. One explanation for the asymmetry is that down-regulated abundances can reach zero, whereas upregulations cannot reach infinity. However, the asymmetry is so pronounced that a biological reason is a strong possibility. Indeed, upregulation of proteins, being caused by protein overexpression, has no natural stopper, while down-regulation is to a large extent driven by protein degradation, which stops or slows down when only strongly bound stoichiometric complexes are left in the system. Thus degradation can provide better correlation between the abundances of the constituent proteins in these complexes.

Entities 2-5 are all represented by Gaussians with very similar widths. Positive and negative components comprise 55% and 33% of total protein pairs, respectively. Entity 2 (“co-down-regulated”) corresponds to protein pairs which have better co-regulation when down-regulated (average r=0.5) than when upregulated (average r=0.3).

Entity 3 (“co-upregulated” proteins) is centered below the diagonal and encompasses proteins correlating better when upregulated (average r=0.5) than when down-regulated (average r=0.28). Entities 2 and 3 account for 23% and 32% of correlating pairs, respectively.

Entity 4 (“anti-correlating”) has a higher anti-correlation when protein A is down-regulated (average r=−0.29 vs r=−0.46), while the above-diagonal entity 5 (“negatively regulated”) shows higher anti-correlation when protein A is upregulated (average r= −0.45 vs r=−0.29). The frequency of protein pairs in these two entities are also almost equal (16% and 17% for entity 4 and 5, respectively).

On the 2D distribution in Fig. 5F, a few protein pairs showed high anti-correlation in one direction and near-zero correlation in the other. For example, in the 50 protein pairs located on the left side of the plot (region 1’), 28 proteins (both “A” and “B” type) mapped to negative regulation of cellular processes (p<0.03). In region 2’, 11 “A” proteins from 50 pairs mapped to “cell cycle” (p<0.005), 5 “B” proteins mapped to “nucleotide excision repair” (p<2.1E-06) and 7 “B” proteins - to “cellular response to DNA damage stimulus” (p<0.022), exposing the former and the two latter pathways as potentially opposing ones (37). The region 4’ mapped onto “ribosome” (16/50 protein pairs, p<2.3E-15) and “ribosome biogenesis” (14 proteins, p<1.0E-12).

Highly anti-correlating pairs of a protein can provide useful information. For example, the upregulated TRIM28 strongly (*r*<-0.61) anti-correlates with 34 proteins. These proteins map to focal adhesion (7 proteins, p<8.1E-06), and a recent study has indeed shown the direct involvement of TRIM28 in cell adhesion (38). Furthermore, the 34 anti-correlating proteins map on ErbB signaling pathway (4 proteins, p<0.001), in line with TRIM28 directly interacting with ErbB4 and suppressing its transcriptional activity (39). Thus, the anti-correlation data can be used in parallel with co-regulation data for deciphering protein function in a given context.

### “Untouchable” proteome reflects essential cell functions

There has been an extensive debate on which house-keeping proteins (HKPs) can be used as steady-level controls in molecular biology experiments (40). Recent research has shown that classic HKPs may actually change their expression under specific conditions (41). For example in our dataset, *GAPDH,* a popular HKP, exhibits stable levels for most treatments, but could not be a good control for studies with bortezomib and few other compounds (Fig. 5J). To identify the best steady-level controls, we investigated the protein expression variations by calculating standard deviations across all perturbations (Fig. 5K and Supplementary Table S11). The typical expression profile of the most steadily expressed and most variable proteins are depicted in Fig. 5L. The 100 most stable proteins belonged to proteasome (9 proteins, p<2E-10), spliceosome (8 proteins, p<3E-05), and mRNA surveillance pathway (6 proteins, p<0.001). On the other hand, the 30 most variable proteins mapped to cell cycle (5 proteins, p<0.001), FoxO signaling pathway (4 proteins, p<0.004) and p53 signaling pathway (3 proteins, p<0.012).

### Increasing the analysis sensitivity by ignoring proximate compounds

Molecules proximate to a compound of interest on the 2/3D maps and/or in hierarchical clusters represent a potential problem for OPLS-DA analysis, as they may share common targets or action mechanism. Contrasting these proximate compounds along with all other molecules against the compound of interest would downplay the common mechanism-related proteins. Therefore, removing proximate compounds from OPLS-DA model would increase the analysis specificity. Indeed, in the model built for docetaxel removing 5 proximate compounds significantly elevated the rankings for tubulin subunits, the known targets of this molecule. When docetaxel was contrasted with all other compounds in the original ProTargetMiner dataset, *TUBB6 TUBA4A TUBA1B TUBB4B TUBB3* and *TUBB* were found on 2^nd^, 8^th^, 15^th^, 25^th^, 46^th^ and 47^th^ positions, while in the reduced model, *TUBB4B TUBA1B TUBB TUBA4A TUBB6 TUBA1A* and *TUBB8* were found on positions 1-7 and *TUBA1C TUBB2A TUBB3* were on 9^th^, 26^th^ and 34^th^ positions, respectively.

### Miniaturizing the ProTargetMiner methodology

In drug development, detailed characterization of compound-induced effects in the most relevant biological setting and cell type is desirable. Thus it would be advantageous to build a dataset similar to ProTargetMiner but with a customized drug panel and cell type. This however, will be both time-consuming and expensive. Miniaturization of the experiment requires determination of the minimal drug panel size for deducing the target and action mechanism. To address this issue, PLS-DA models were built in R by including different numbers of contrasting compounds in the model (*n* = 1-54, 50 molecule combinations randomized for each *n*). The mean rankings of the known drug targets for representative compounds camptothecin, methotrexate, OSW-1 and paclitaxel were determined for each number of contrasting drugs *n*. As expected, with higher *n* the deconvolution process was more successful for drug targets but not for random proteins (Fig. 6). Encouragingly, already 8-10 molecules in the drug panel were in most cases enough for target rankings below 10.

**Figure 6.**
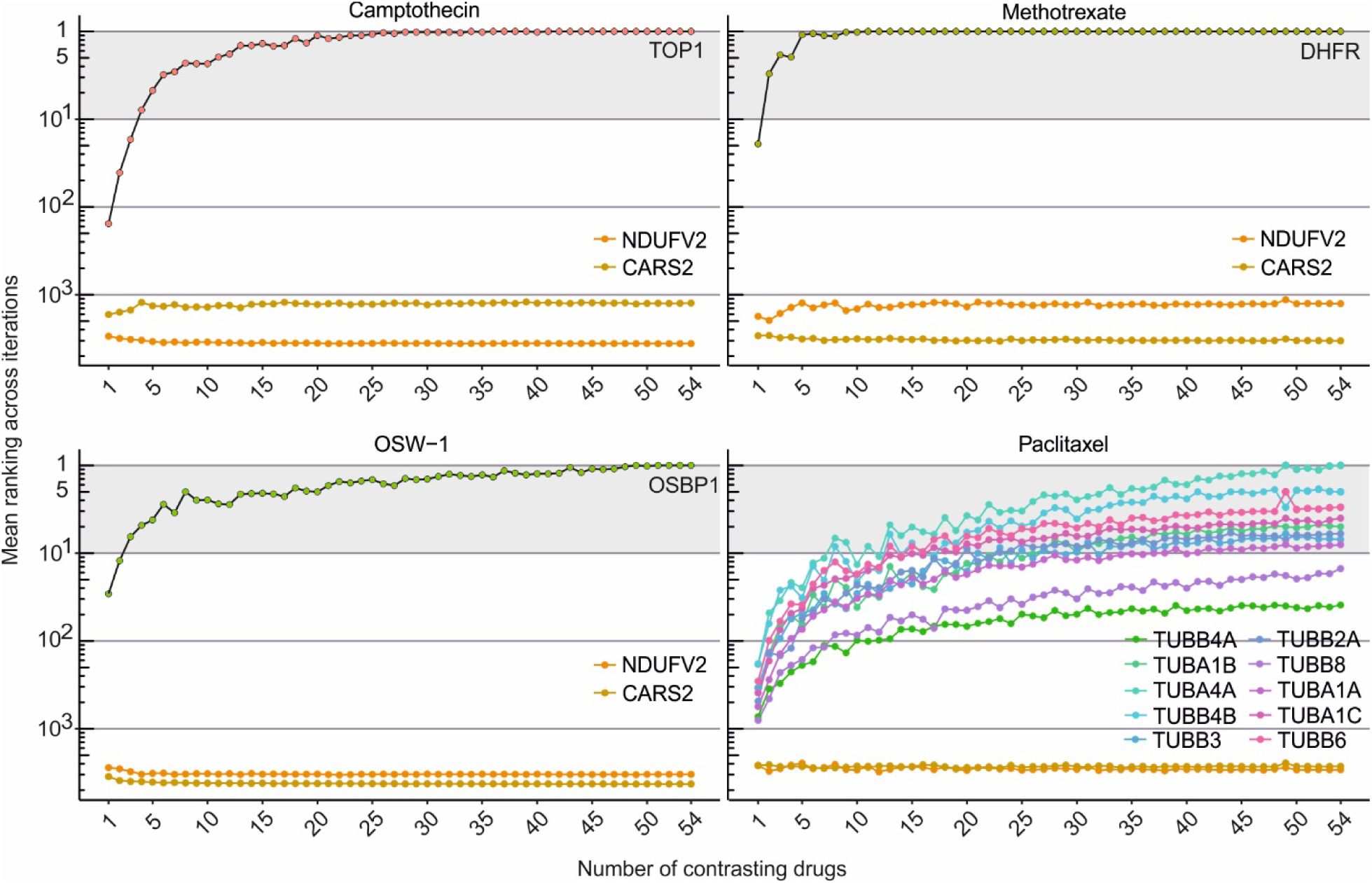
Miniaturization of ProTargetMiner. Determination of the optimal number of contrasting molecules for efficient target deconvolution by FITExP using ProTargetMiner data. Four compounds were contrasted against 50 random combinations of 1-54 compounds by PLS modeling and the mean ranking was calculated for each number. *NDUFV2* and *CARS2* were randomly chosen.

### Deep proteome validation analysis

A repeated, deep proteome analysis was performed to confirm the consistency of the deduced drug targets and action mechanisms. To provide maximum diversity of the proteome responses, each of the 9 drugs used in this experiment was chosen from a separate cluster in Fig. 2. In total, 94,061 peptides were quantified belonging to 8,898 proteins, of which 8,308 proteins were defined by at least two peptides. After removing the missing values, 7,398 proteins were used for analysis (Supplementary Table S12). Interestingly, PCA showed 10 principal components, explaining from 28% of the data for the 1^st^ component to 2.7% for the last component (Fig. 7A-B). This result confirmed the orthogonality of the death trajectories induced by the chosen drugs.

**Figure 7.**
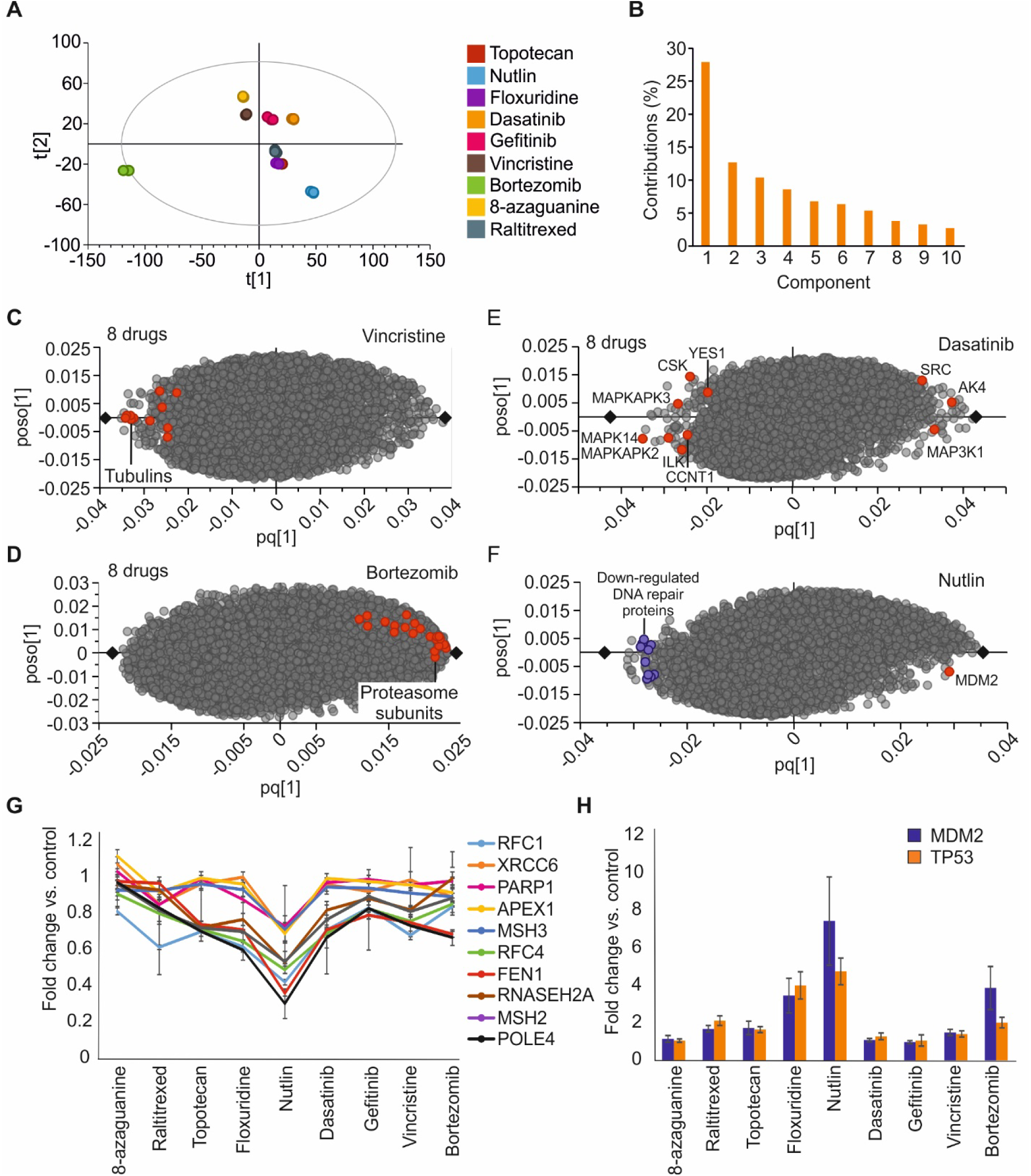
A deep proteomics validation set confirmed the ProTargetMiner findings. **A,** OPLS plot showing that compound signatures can be reproducibly separated (7398 proteins). **B,** The existence of 10 orthogonal dimensions in PCA data and the contribution of each component. **C,** OPLS-DA plots for bortezomib (**C**), and vincristine (**D**) showing the specific upregulation of proteasome subunits for bortezomib and down-regulation of tubulin subunits for vincristine. **E,** OPLS-DA modeling revealed 5 known targets among top proteins for dasatinib. **F,** The OPLS-DA model contrasting nultin against 8 drugs. The highlighted DNA repair proteins were specifically down-regulated. **G,** DNA repair proteins identified in the OPLS-DA model showed a specific dip in expression for nutlin vs other compounds. **H,** Plotting the expression of *MDM2* and p53 showed that these proteins are most specific to nutlin. Data are represented as mean±SD.

The OPLS-DA models built for bortezomib and vincristine (Fig. 7C-D) yielded nearly identical target/mechanistic proteins as the models built using the main dataset (Fig. 3). In an OPLS model for dasatinib, 10 known targets (among 23 known targets in DrugBank derived from different studies) were confirmed (Fig. 7E). Importantly, at least five kinases were among the most specifically changed proteins, with MAPK14 being the most specifically down-regulated protein. These findings show that ProTargetMiner methodology can be successfully miniaturized and that the scheme can be applicable to kinase inhibitors in the state-of-the-art proteomics depth.

### Specificity helps to identify subtle but biologically significant changes

The case of nutlin illustrates the importance of using the specificity parameter as opposed to conventional differential regulation in determining drug mechanism. Nutlin has been found to slow down the DNA repair, probably through *MDM2* mediated stalling of double strand break repair (42). While the 30 most down-regulated proteins by nutlin mapped to “cell cycle” (20 proteins, p<2E-15), the 30 most *specifically* down-regulated proteins extracted from the OPLS-DA model (Fig. 7F) gave “DNA repair” (9 proteins, p<2E-5). The regulation of these proteins across the drug panel shows a dip for nultin (Fig. 7G). The top DNA repair related protein *APEX1* ranked 10^th^ among the most *specifically* down-regulated proteins out of the total 7398 proteins in the OPLS model, while it was the 520^th^ most down-regulated protein, exhibiting only a −41% abundance change. As the majority of the DNA repair proteins were only marginally regulated by nutlin, these proteins would be difficult if not impossible to attribute to the action mechanism without contrasting nutlin against other compounds. Also, both p53 and its negative regulator *MDM2*, the known nutlin target (43), exhibited higher specific response to nultin than to other compounds (Fig. 7H). These results unravel a potential approach to study subtle but biologically meaningful changes.

## Discussion

We generated the first installment of a proteomics signature library for anticancer molecules, and proposed a modeling scheme that could be employed for deconvolution of drug targets, action mechanism, resistance factors, overall cellular effects and more. We consider ProTargetMiner as a complement for preceding drug deconvolution databases, such as CMap (2,3). The OPLS-DA modeling used in ProTargetMiner can be hypothetically applied to transcriptomics data or to non-anti-cancer treatments, as long as the biological endpoints for all used compounds are similar, as in ProTargetMiner. We limited our discussion to cases in which a single drug is contrasted against many others, but the same approach can also be applied to characterize features shared among a selected class of compounds, or between any combinations of drugs. Such a methodology can also be applied to a panel of cell lines, *e.g.*, inquiring which proteins specifically respond to a compound in a given cell line but not in other cell types.

The 11 dimensions discovered in the ProTargetMiner dataset by factor analysis represent orthogonal pathways, theoretical constructs that do not necessarily resemble the classical pathways known to biologists from textbooks (44). The 2012 report of the Cell Death Nomenclature Committee recognized 13 distinct death modalities, while the 2018 updated report recognized 14 death modalities. These numbers are not far from the dimensionality we discovered. The orthogonal pathways uncovered here deserve a detailed bioinformatics analysis, as they may hide novel death modalities.

Our analysis showed clearly that performing simple differential expression and focusing on the top hits obscures subtle but specific compound-induced regulations. These levels of regulation are only accessible to high precision technologies. In proteomics, precision comes not only from the instrumental parameters, but also from post-processing such as factor analysis (45). We showed that specific regulations as small as 15% can be reliably assessed as the most characteristic changes for a given compound. Such level of precision would have been impossible several years ago, when the standard practice in proteomics recommended disregarding the regulations less than a factor of 1.5 or even 2. Such subtle but specific regulations could be approached here using the specificity concept through building OPLS-DA models that automatically take into account reproducibility, missing values and the change magnitude.

ProTargetMiner can be used for a variety of exploratory studies, only a few of which we have so far performed. Among yet to be explored opportunities is the discovery of novel biological pathways or gene function elucidation at the protein level. These studies can be done based on the proteomics co-regulation database. Our analysis also revealed a host of anti-correlating protein pairs, which can be biologically meaningful. An unexpected discovery was the number of anti-correlations (spots 4 & 5 on Fig. 5F), reaching 3/5 of that of positive correlations (spots 2&3). Being largely ignored currently in bioinformatics analysis, these anti-correlations represent an untapped wealth of important biological information. Furthermore, the dataset allows one to identify proteins that can serve as HKPs in molecular biology experiments.

This resource is expandable in various dimensions, *e.g.*, by incorporating more perturbations, time points, and profiling more cell lines. Although expansion of the compound library seems desirable, one must consider that for a comprehensive database, enough perturbations must be done to saturate all possible cellular states. Ideally, highly specific inhibitors of every cellular protein are required. But given the astronomical number of potential perturbants, building a truly comprehensive library is an open-end project.

While precision medicine targets specific cell types with defined mutations, building a comprehensive proteome response database for every such cell type is impossible. Fortunately, as we have shown, ProTargetMiner approach can be easily customized and miniaturized. With top-of-the-line proteomics instruments reaching the depth of ≥10,000 proteins (46), a triplicate analysis of 9 perturbations can be performed in less than a week.

The steady-expression part of the proteome represents one of the biggest puzzles uncovered in this work. Would targeting these proteins result in inevitable cell demise? In this case, this “untouchable” proteome may represent a collection of potential antiproliferation targets. As of April 2018, only PSMB1 and KHSRP are indicated as the target of FDA approved anticancer drugs in DrugBank among the top 50 untouchable proteins, while PSMA1 and 5 as well as SF3A3 are targets for experimental compounds. How many more potential targets does this enigmatic group of proteins hide?

Summarizing, ProTargetMiner is a novel tool with significant potential in drug discovery, which can be useful to broad community of cancer researchers.

## Acknowledgements

We would like to acknowledge Marie Ståhlberg and Carina Palmberg for their assistance in mass spectrometry analyses. We are also grateful to Elias S.J. Arnér for providing TRi-1 and TRi-2 compounds, Stig Linder for b-AP15 and Kaori Sakurai for OSW-1 and tomatine.

## Materials and Methods

### Compounds

The initial drug library consisted of 118 molecules cherry-picked for cancer from Selleckchem FDA-approved drug library. AXL inhibitor TP-0903 was bought from Selleckchem (Cat#S7846) and auranofin from Sigma (Cat#A6733).

### A549 Cell culture

Human A549 cells (Sigma, USA; RRID:CVCL_0023; Cat#86012804), established from lung carcinomatous tissue from a 58-year-old Caucasian male, were obtained in September 2015. Low passage numbers (n<10) were used in all the experiments from frozen aliquots of the same source (passage number 2). Cells were routinely tested for presence of mycoplasma every month using the MycoAlert Mycoplasma Detection Kit (Fischer Scientific; Cat#11650261). Cells were grown in DMEM medium (Fisher Scientific; Cat#11625200) supplemented with 10% FBS (Fisher Scientific; Cat#11560636), 2 mM L-glutamine (Fisher Scientific; Cat#BE17-605E) and 100 units/mL of penicillin/streptomycin (Thermo Fisher; Cat#15140122) and incubated at 37°C in 5% CO_2_. In LC50 determination, cells were seeded at a density of 4000/well in 96 well plates and after a day of growth, treated with the molecules for 48 h. Thereafter cell viability was measured using CellTiter-Blue® Cell Viability Assay (Promega; Cat#G8081) according to the manufacturer protocol.

### Proteomics

For proteomics analysis, the cells were seeded at a density of 250k per well and allowed to grow for 24 h in biological triplicates. Next, cells were either treated with vehicle (DMSO) or compounds at LC50 concentrations. Each 10 experiments included one vehicle-treated control, 3 control drugs and 6 library compounds. After treatment, cells were collected, washed twice with PBS (Fisher Scientific; Cat#11629980) and then lysed using 8 M urea (Sigma; Cat#U5378), 1% SDS, and 50 mM Tris at pH 8.5 with protease inhibitors (Sigma; Cat #05892791001). The cell lysates were subjected to 1 min sonication on ice using Branson probe sonicator and 3 s on/off pulses with a 30% amplitude. Protein concentration was then measured for each sample using a BCA Protein Assay Kit (Thermo; Cat#23227). 50 µg of each sample was reduced with DTT (final concentration 10 mM) (Sigma; Cat#D0632) for 1 h at room temperature. Afterwards, iodoacetamide (IAA) (Sigma; Cat#I6125) was added to a final concentration of 50 mM. The samples were incubated in room temperature for 1 h in the dark, with the reaction being stopped by addition of 10 mM DTT. After precipitation of proteins using methanol/chloroform, the semi-dry protein pellet was dissolved in 25 µL of 8 M urea in 20 mM EPPS (pH 8.5) (Sigma; Cat#E9502) and was then diluted with EPPS buffer to reduce urea concentration to 4 M. Lysyl endopeptidase (LysC) (Wako; Cat#125-05061) was added at a 1 : 100 w/w ratio to protein and incubated at room temperature overnight. After diluting urea to 1 M, trypsin (Promega; Cat#V5111) was added at the ratio of 1 : 100 w/w and the samples were incubated for 6 h at room temperature. Acetonitrile (Fisher Scientific; Cat#1079-9704) was added to a final concentration of 20% v/v.

TMT10 reagents (Thermo; Cat#90110) were added 4x by weight to each sample, followed by incubation for 2 h at room temperature. The reaction was quenched by addition of 0.5% hydroxylamine (Thermo Fisher; Cat#90115). Samples were combined, acidified by trifluoroacetic acid (TFA; Sigma; Cat#302031-M), cleaned using Sep-Pak (Waters; Cat#WAT054960) and dried using a DNA 120 SpeedVac™ concentrator (Thermo).

Samples were then resuspended in 0.1% TFA, and separated into 8 fractions using High pH Reversed-Phase Peptide Fractionation Kit (Thermo; Cat#84868). After resuspension in 0.1% FA (Fisher Scientific), each fraction was analyzed with a 90 min gradient in randomized order.

The deep proteome validation samples (tags assigned in Supplementary Table S2) were prepared according to the above protocol until the multiplexing, cleaning and drying steps, after which the samples were resuspended in 20 mM ammonium hydroxide and separated into 96 fractions on an XBrigde BEH C18 2.1×150 mm column (Waters; Cat#186003023), using a Dionex Ultimate 3000 2DLC system (Thermo Scientific) over a 48 min gradient of 1-63%B (B=20 mM ammonium hydroxide in acetonitrile) in three steps (1-23.5%B in 42 min, 23.5-54%B in 4 min and then 54-63%B in 2 min) at 200 µL/min flow. Fractions were then concatenated into 16 samples in sequential order (*e.g.* 1,17,33,49,65,81). After drying and resuspension in 0.1% formic acid (FA) (Fisher Scientific), each fraction was analyzed with a 90 min gradient (total method time = 110 min) in random order.

### LC-MS analysis

Samples were loaded with buffer A (0.1% FA in water) onto a 50 cm EASY-Spray column (75 µm internal diameter, packed with PepMap C18, 2 µm beads, 100 Å pore size; Cat#ES803) connected to the EASY-nLC 1000 (Thermo; Cat#LC120) and eluted with a buffer B (98% ACN, 0.1% FA, 2% H_2_O) gradient from 2% to 35% of at a flow rate of 250 nL/min. Mass spectra were acquired with an Orbitrap Q Exactive Plus mass spectrometer (Thermo; Cat# IQLAAEGAAPFALGMBDK) in the data-dependent mode at a nominal resolution of 30,000, in the m/z range from 375 to 1400. Peptide fragmentation was performed via higher-energy collision dissociation (HCD) with energy set at 35 NCE.

For deep proteomics validation set, samples were loaded with buffer A (0.1% FA in water) onto a 50 cm EASY-Spray column (75 µm internal diameter, packed with PepMap C18, 2 µm beads, 100 Å pore size) connected to a nanoflow Dionex UltiMate 3000 UPLC system (Thermo) and eluted in an increasing organic solvent gradient from 2% to 26% (B: 98% ACN, 0.1% FA, 2% H_2_O) at a flow rate of 300 nL/min. Mass spectra were acquired with a Q Exactive HF mass spectrometer (Thermo; Cat#IQLAAEGAAPFALGMBFZ) in data-dependent mode at a nominal resolution of 60,000 (@200 *m/z*), in the mass range from 350 to 1500 *m/z*. Peptide fragmentation was performed via higher-energy collision dissociation (HCD) with energy set at 33 NCE.

### Protein identification and quantification

The raw data from LC-MS were analyzed by MaxQuant, version 1.5.6.5 (RRID:SCR_014485) (47). The Andromeda engine searched MS/MS data against Uniprot complete proteome database (human, version UP000005640_9606, 92957 entries). Cysteine carbamidomethylation was used as a fixed modification, while methionine oxidation was selected as a variable modification. Trypsin/P was selected as enzyme specificity. No more than two missed cleavages were allowed. A 1% false discovery rate was used as a filter at both protein and peptide levels. For all other parameters, the default settings were used.

### Statistical analysis

After removing all the contaminants, only proteins with at least two peptides were included in the final dataset. Protein abundances were normalized by the total protein abundance in each sample. Data were processed by Excel, R, Python, and SIMCA (Version 15, UMetrics, Sweden; RRID:SCR_014688).

### Protein correlation analysis

Relative protein abundances in log2-scale were reported by Diffacto (45) and subject to batch effect removal using the Limma package (48). We excluded 345 proteins that were quantified in less than 30 individual LC-MS/MS runs, and averaged the protein abundances from replicate experiments for each individual drug treatment. Afterwards, a 4212-by-4212 correlation matrix was calculated. Two-tail p-values associated with the Pearson’s correlation coefficients were calculated based on t-distribution, and were subjected to the Benjamin-Hochberg procedure for controlling false discovery rate (FDR). A cutoff of FDR-adjusted p-value < 0.001 was applied to select strongly correlating or anti-correlation protein pairs. Considering the correction of in total 17,740,944 (4212−4212) p-values, the FDR cutoff is rather stringent

### Network mapping

STRING version 10.5 (http://string-db.org) tool was employed for protein network analysis (49). Medium confidence threshold (0.4) was used to define protein-protein interactions. Enrichment analysis with the whole genome as a background dataset was used to identify the enriched gene ontology terms and pathways.

### Data availability

The LC-MS/MS raw data files are deposited in the jPOST repository of the ProteomeXchange Consortium under the dataset identifier PXD009775 (original ProTargetMiner data) and PXD009644 (deep proteomics validation set) (50). The extracted protein abundances data and relevant outputs of data analysis are provided in supplementary tables cited in the text.

## Supplemental Figures

**Figure S1.**
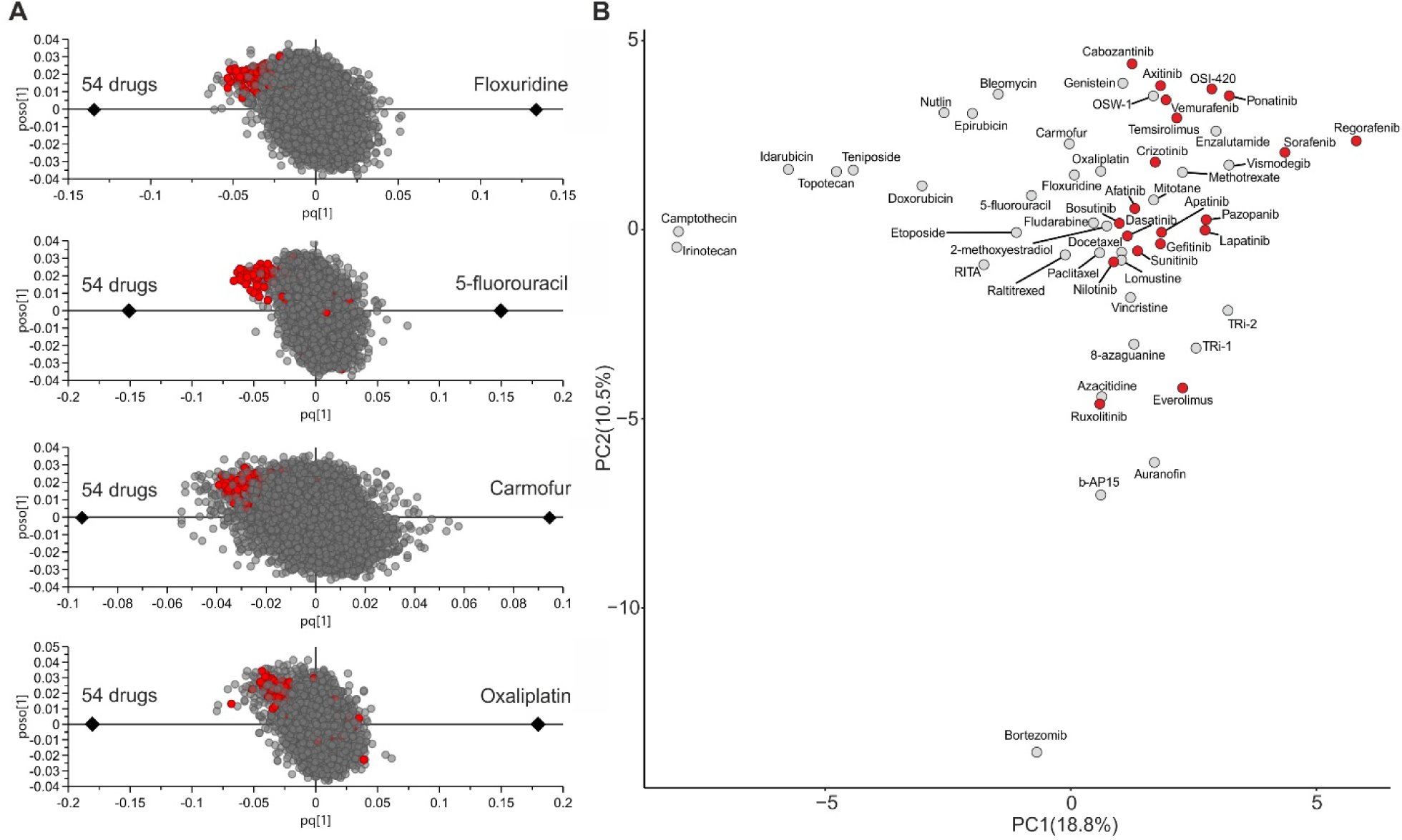
ProTargetMiner reveals compound effect on protein complexes and might be applicable to kinase inhibitors. **A,** Pyrimidine analogues lead to ribosomal biogenesis stress and cell death. Oxaliplatin, an alkylating agent produced a similar effect. **B,** PCA analysis demonstrating that the regulation data of 68 kinases can separate kinase inhibitors from the rest of the library to some extent, especially from proteasome and topoisomerase inhibitors.

**Figure S2.**
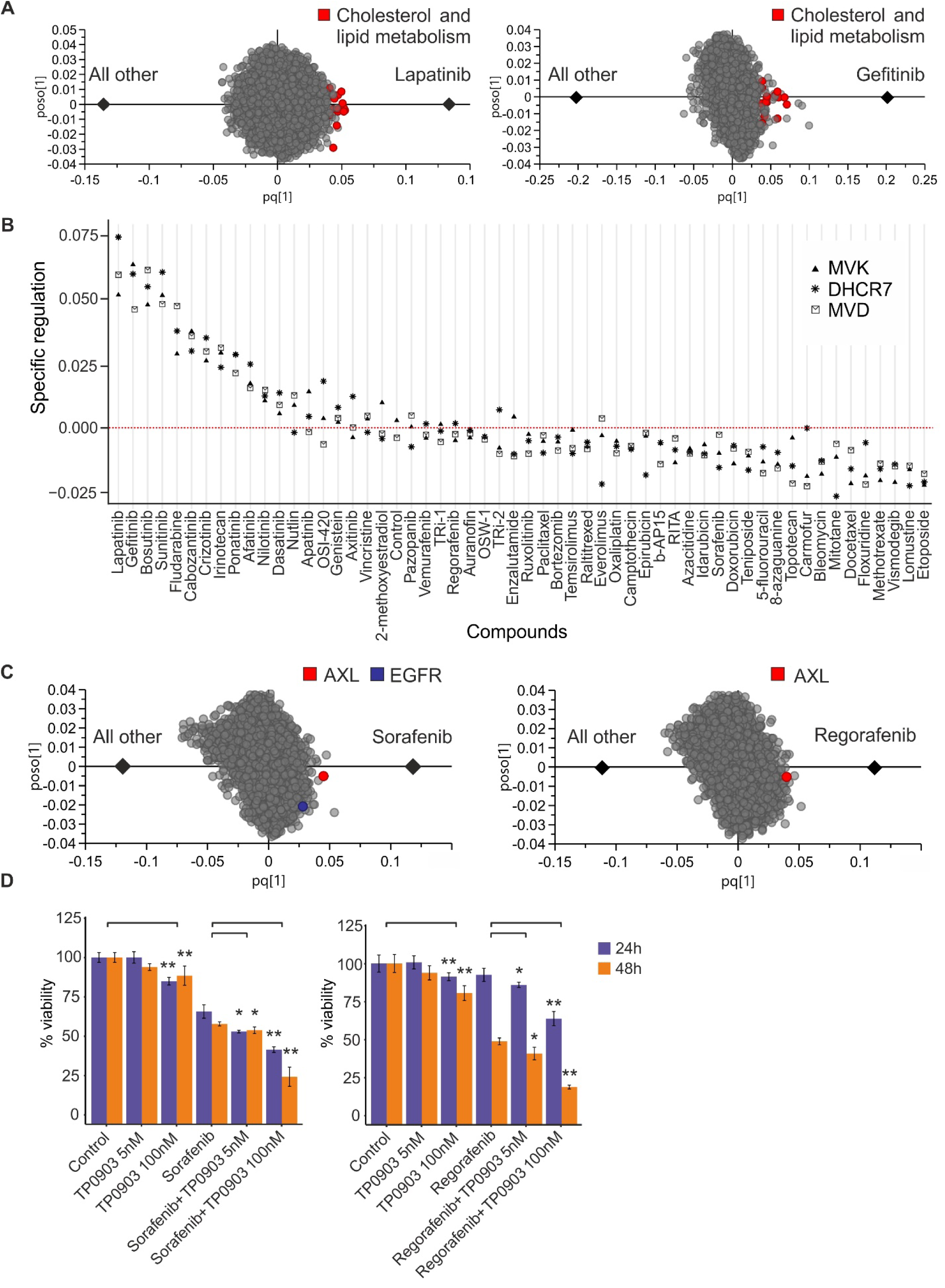
ProTargetMiner yields molecular information for kinase inhibitors that could be mechanistically relevant. **A,** Upregulation of proteins involved in cholesterol and lipid metabolism in response to lapatinib and gefitinib. **B,** Representative examples of such upregulated proteins in the panel of drugs demonstrate that several kinase inhibitors upregulate proteins in these pathways. **C,** Models contrasting sorafenib and regorafenib against 44 other compounds, showing the specific upregulation of AXL upon treatment of A549 cells (note the upregulation of EGFR by sorafenib). **D,** Treatment of cells with non-cytotoxic concentrations of TP0903, a specific and nanomolar AXL inhibitor, sensitized A549 cells to the effect of sorafenib and regorafenib in 24 and 48h (*p<0.05, **p<0.005). Data are represented as mean±SD.

**Figure S3.**
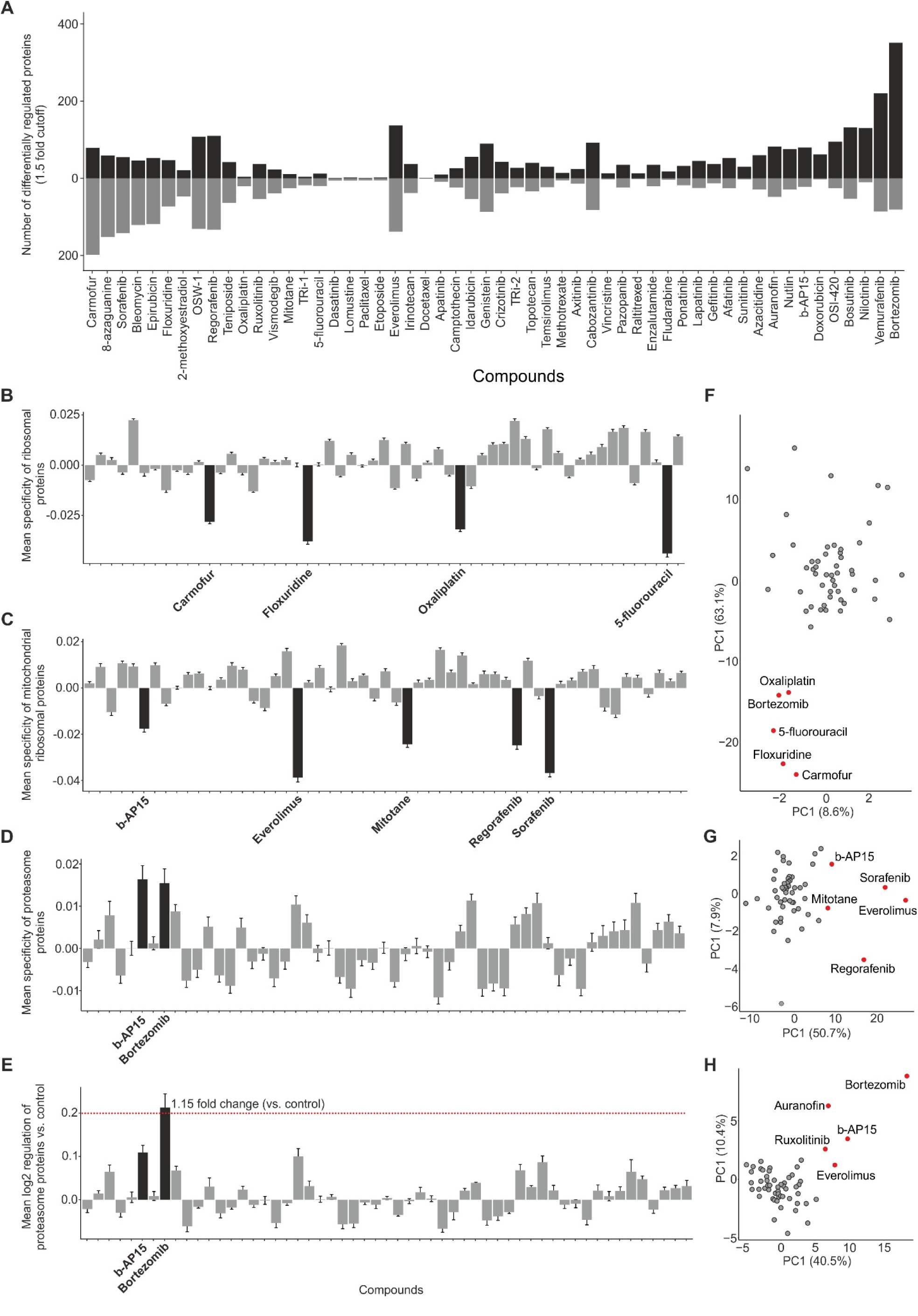
The extent of proteome deviation induced by each compound and complex-specific effects of compounds. **A,** Simple calculation of the number of differentially regulated proteins shows the proteo-active compounds and the extent of proteome change for each molecule **B,** Mean specific regulation of 74 ribosomal proteins for compound panel, showing the effect of the highlighted compounds on ribosome biogenesis. **C,** Mean specific regulation of 66 mitochondrial ribosomal proteins for compound panel, showing the effect of highlighted compounds on this complex. **D,** Mean specific regulation of 40 proteasome subunits for the panel of compounds. Proteasome inhibitors have been highlighted. **E,** The average regulation of proteasome subunits in the panel of compounds, demonstrating that ProTargetMiner can detect effects as subtle as 15% fold change (the effect is even mitigated due to inclusion of all proteasome subunits). PCA analyses of compounds based on (**F**) 74 ribosomal proteins, (**G**) 57 mitochondrial ribosomal proteins and (**H**) 38 proteasome subunits with no missing values in the ProTargetMiner dataset. The analysis clearly shows the effect of selected compounds on these complexes and indicates that specific proteome subsets are useful in differentiating compound effects. Data are represented as mean±SD.

**Figure S4.**
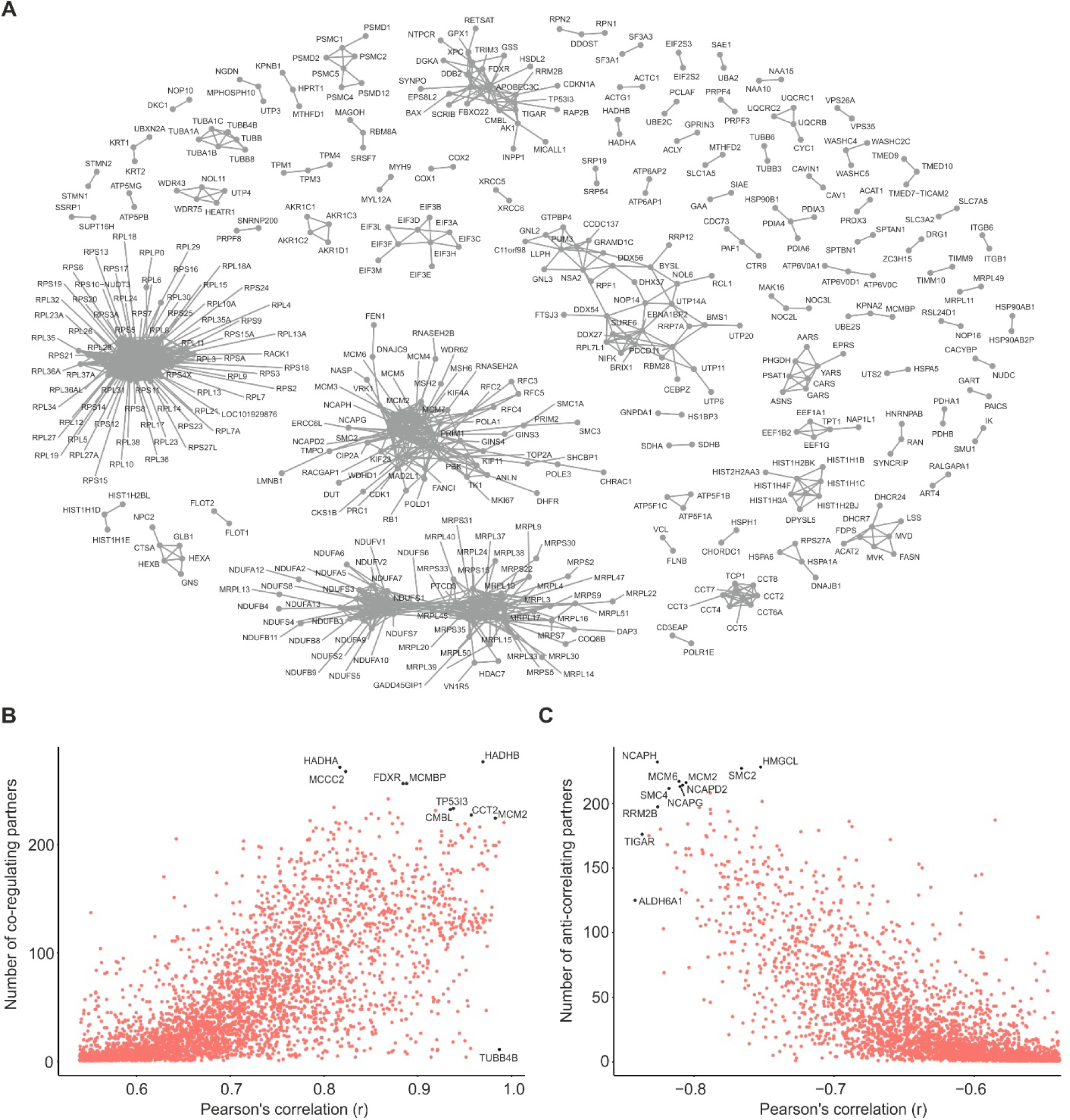
ProTargetMiner co-regulation database can pull out functionally coherent groups of genes. **A,** The association of top 2500 co-regulated pairs (many were redundant due to presence in the same complexes), mostly representing proteins from the same complexes or pathways. The number of (**B**) co-regulating and (**C**) anti-correlating partners for each protein vs and their highest correlation.

## Supplemental tables

**Table S1.** Compounds, curated targets from DrugBank and class assignments.

| Drug name | Targets | Class |
| --- | --- | --- |
| Axitinib | FLT1, FLT4, KDR | Kinase inhibitor |
| Lapatinib | ERBB2, EGFR | Kinase inhibitor |
| Crizotinib | ALK, MET | Kinase inhibitor |
| Regorafenib | FLT1, KDR, FLT4, KIT, PDGFRA, PDGFRB, FGFR1, FGFR2, DDR2, TEK, NTRK1, RAF1, EPHA2, MAPK11, BRAF, FRK, ABL1, RET | Kinase inhibitor |
| Ruxolitinib | JAK1, JAK2 | Kinase inhibitor |
| Afatinib | ERBB2, ERBB4, EGFR | Kinase inhibitor |
| Cabozantinib | MET, RET, KDR | kinase inhibitor |
| Ponatinib | BCR, ABL1, KIT, RET, TEK, FLT3, FGFR1, FGFR2, FGFR3, FGFR4, LCK, SRC, LYN, KDR, PDGFRA | kinase inhibitor |
| Sorafenib | RAF1, BRAF, FLT4, KDR, FLT3, PDGFRB, KIT, FGFR1, RET, FLT1, | kinase inhibitor |
| Bosutinib | LYN, BCR, ABL1, HCK, SRC, CDK2, MAP2K1, MAP3K2, MAP2K2, CAMK2G | kinase inhibitor |
| Sunitinib | PDGFRB, FLT1, KIT, KDR, CSF1R, FLT4, FLT3, PDGFRA | kinase inhibitor |
| Dasatinib | ABL1, SRC, EPHA2, LCK, YES1, KIT, PDGFRB, STAT5B, ABL2, FYN, BTK, NR4A3, BCR, CSK, EPHA5, EPHB4, FGR, FRK, HSPA8, LYN, ZAK, MAPK14, PPAT | kinase inhibitor |
| Apatinib | KDR (VEGFR2) | kinase inhibitor |
| Gefitinib | EGFR | kinase inhibitor |
| Vemurafenib | BRAF | kinase inhibitor |
| OSI-420 | EGFR | kinase inhibitor |
| Nilotinib | KIT, ABL1 | Kinase inhibitor |
| Pazopanib | FLT1, KDR, PDGFRA, PDGFRB, FLT4, FGFR3, KIT, FGF1, ITK, SH2B3 | Kinase inhibitor |
| Vismodegib | SMO | Hedgehog signaling pathway |
| Azacitidine | DNMT1, nucleotide | DNA methyltransferase inhibitor |
| Everolimus | mTOR | mTOR inhibitor |
| Temsirolimus | mTOR | mTOR inhibitor |
| Fludarabine | POLA1, RRM1, DCK, nucleotide | Antimetabolite |
| Oxaliplatin | Nucleotide | Antimetabolite |
| 5-fluorouracil | TYMS, Nucleotide | Antimetabolite |
| Methotrexate | DHFR | Antimetabolite |
| Raltitrexed | FPGS, TYMS | Antimetabolite |
| Carmofur | TYMS and AC (acid ceramidase) inhibitor | Antimetabolite |
| Floxuridine | TYMS | Antimetabolite |
| 8-azaguanine | PNP | Antimetabolite |
| Vincristine | TUBB, TUBA4A | Tubulin polymerization inhibitors |
| 2-methoxyestradiol | HIF1A, COMT, CYP1A1, CYP1B1, CYP19A1 | Tubulin polymerization inhibitors |
| Paclitaxel | BCL2, TUBB1, NR1I2, MAP4, MAP2, MAPT | Tubulin depolymerization inhibitors |
| Docetaxel | TUBB1, bcl2, MAP2, MAP4, MAPT, NR1I2 | Tubulin depolymerization inhibitors |
| Genistein | ESR2, ESR1, TOP2A, PTK2B, NCOA2, NCOA1 | Topoisomerase II inhibitor, protein kinase inhibitor |
| Epirubicin | CHD1, TOP2A, nucleotide | Topoisomerase II inhibitor, DNA intercalation |
| Doxorubicin | TOP2A, Nucleotide | Topoisomerase II inhibitor, DNA alkylating |
| Etoposide | TOP2A, TOP2B | Topoisomerase II inhibitor |
| Idarubicin | TOP2A, Nucleotide | Topoisomerase II inhibitor, DNA alkylating |
| Lomustine | STMN4, nucleotide | Alkylating agent |
| Teniposide | TOP2A | Topoisomerase II inhibitor |
| Topotecan | TOP1MT, TOP1, nucleotide | Topoisomerase I inhibitor |
| Irinotecan | TOP1, TOP1MT | Topoisomerase I inhibitor |
| Camptothecin | TOP1 | Topoisomerase I inhibitor |
| Mitotane | CYP11B1, FDX1 | Unknown, peripheral metabolism of steroids and adrenocortical steroid inhibitor |
| RITA | MDM2 | MDM2 inhibitor |
| Nutlin | MDM2-p53 interaction | MDM2 inhibitor |
| Enzalutamide | AR (androgen receptor) | Androgen receptor inhibitor |
| Bleomycin | LIG1, LIG3, nucleotide | Induction of DNA strand breaks |
| Bortezomib | PSMB5, PSMB1 | Proteasome inhibitor |
| b-AP15 | UCL5, USP14 | Proteasome deubiquitinase inhibitor |
| Auranofin | TrxR1, PRDX5, IKBKB, (USP14 and UCL5 inhibition has also been seen) | Thioredoxin reductase 1 inhibitor, proteasome deubiquitinase inhibitor |
| TRI-1 | TrxR1 | Thioredoxin reductase 1 inhibitor |
| TRI-2 | TrxR1 | Thioredoxin reductase 1 inhibitor |
| Tomatine | Acetylcholinesterase, membrane disruption | Unknown, disruption of membrane, potential inhibition of the proteasome |
| OSW-1 | OSBP1 | Unknown, OSBP binder |

**Table S2.** TMT-10 multiplexing information for the ProTargetMiner experiments.

|  | Experiment 1 | Experiment 2 | Experiment 3 | Experiment 4 | Experiment 5 | Experiment 6 | Experiment 7 | Experiment 8 | Experiment 9 | Deep validation experiment |
| --- | --- | --- | --- | --- | --- | --- | --- | --- | --- | --- |
| 126 | 5-fluorouracil | Regorafenib | Vismodegib | Cabozantinib | Fludarabine | - | Idarubicin | Carmofur | OSW-1 | Control |
| 127N | Methotrexate | Methotrexate | Methotrexate | Methotrexate | Methotrexate | Methotrexate | Methotrexate | Methotrexate | Methotrexate | 8-azaguanine |
| 127C | Paclitaxel | Paclitaxel | Paclitaxel | Paclitaxel | Paclitaxel | Paclitaxel | Paclitaxel | Paclitaxel | Paclitaxel | Raltitrexed |
| 128N | Doxorubicin | Afatinib | Axitinib | Ponatinib | Sunitinib | Etoposide | Temsirolimus | Genistein | Auranofin | Topotecan |
| 128C | Irinotecan | Bortezomib | Nilotinib | Everolimus | Dasatinib | Gefitinib | Enzalutamide | Bleomycin | TRI-1 | Floxuridine |
| 129N | Camptothecin | Camptothecin | Camptothecin | Camptothecin | Camptothecin | Camptothecin | Camptothecin | Camptothecin | Camptothecin | Nutlin |
| 129C | RITA | Ruxolitinib | Crizotinib | Sorafenib | Docetaxel | Oxaliplatin | Vemuratenib | 8-Azaguanine | TRI-2 | Dasatinib |
| 130N | Nutlin | Azacitidine | Topotecan | Bosutinib | Lomustine | Mitotane | OSI-420 | Floxuridine | b-AP15 | Gefitinib |
| 130C | Raltitrexed | Pazopanib | Lapatinib | Epirubicin | Vincristine | Apatinib | Teniposide | 2-methoxyestra diol | Tomatine | Vincristine |
| 131 | Control | Control | Control | Control | Control | Control | Control | Control | Control | Bortezomib |

**Table S5.**
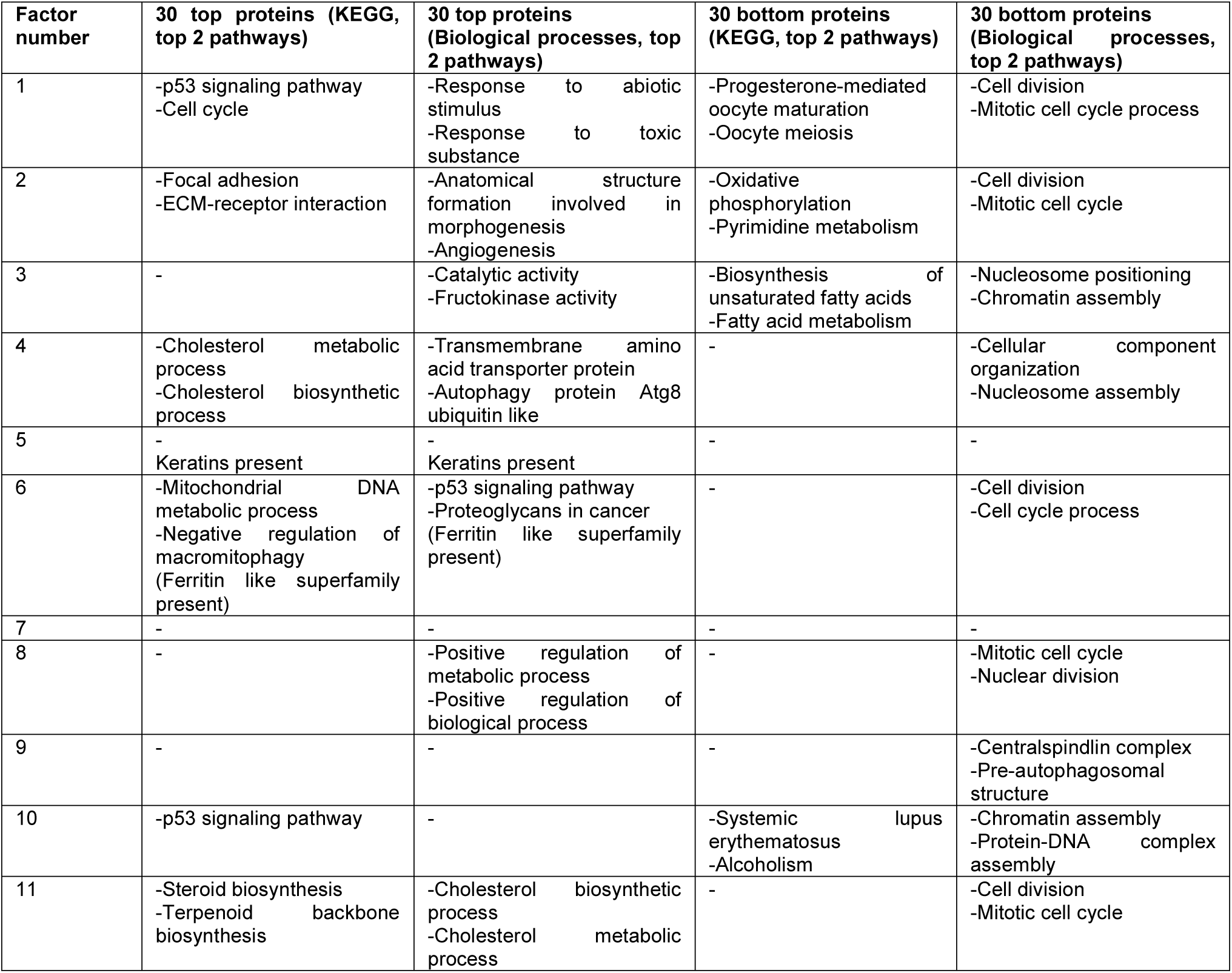
The pathway analysis of most contributing proteins to 11 dimensions in the ProTargetMiner dataset. Top proteins contribute when up-regulated. Bottom proteins contribute when down-regulated.

### Supplementary Table legends

**Table S1.** Compounds, curated targets from DrugBank and class assignments.

**Table S2.** TMT-10 multiplexing information for the ProTargetMiner experiments.

**Table S3.** The original ProTargetMiner dataset with 56 drugs.

**Table S4.** The contribution of all proteins to the 11 dimensions found in the ProTargetMiner dataset using Factor analysis.

**Table S5.** The pathway analysis of most contributing proteins to 11 dimensions in the ProTargetMiner dataset. Top proteins contribute when up-regulated. Bottom proteins contribute when down-regulated.

**Table S6.** The specificity of each protein in response to each compound against all other compounds in OPLS models.

**Table S7.** The constituent proteins and enriched pathways in the 4212 x 4212 protein-protein correlation matrix.

**Table S8.** The co-regulated and anti-correlating protein pairs below FDR<0.001 across the perturbations in ProTargetMiner. The second tab represents proteins with no strong correlation with any other protein.

**Table S9.** The external payload data (nodes as well as positive and negative edges) for ProTargetMiner co-regulation and anti-correlation database which can be uploaded to the StringDB.

**Table S10.** Protein A vs. B correlations calculated separately for above and below the median abundance of protein A, in ProTargetMiner.

**Table S11.** Standard deviations of protein expression across all perturbations in ProTargetMiner (untouchable and variable proteomes).

**Table S12.** The deep proteomics validation dataset with 9 drugs.

